# miR-190 is a Key Regulator in Establishing Cell Polarity and Specification in the Drosophila Nervous System

**DOI:** 10.1101/2025.05.28.656688

**Authors:** Gerson Ascencio, Laura Galvan, Julie M. Sanchez, Ariana Nagainis, Cynnie Tam, Lauren M. Goins, Blake Riggs

## Abstract

Asymmetric cell division generates cellular diversity in developing tissues, particularly in the CNS. In *Drosophila* neuroblasts, this process relies on polarity complexes and fate determinants, yet its molecular regulation remains unclear. Here, we identify miRNA-190 as a key regulator of neuroblast polarity and differentiation. Single-cell RNA sequencing and transcriptome analysis reveal that miR-190 deficiency disrupts CNS cell populations, reducing neurons while increasing neural progenitors and glia. Mechanistically, miR-190 is required for proper localization of the Par complex and basal determinants during mitosis. In miR-190 mutants, these factors mislocalize, leading to defective polarity and fate specification in embryonic neuroblast. qPCR analysis shows that miR-190 targets *RhoGAP*, which modulates Cdc42 activation and Par-6, crucial factors in neuroblast polarity. We propose a model in which miR-190 ensures proper Cdc42 activation and polarity establishment by targeting transcripts for degradation. miR-190 has been implicated in various cancers, and our findings provide a mechanistic framework for understanding miR-190’s roles in tumorigenesis and its broader involvement in metabolic diseases.

## Introduction

The generation of cell diversity is accomplished by an asymmetric partitioning of cell fate determinants during cell division. This asymmetric cell division (ACD) is an evolutionarily conserved mechanism used by multicellular organisms to control cell fate and generate cell diversity by partitioning factors that either establish a differentiation state or a self-renewal state. The neural progenitor cells in the development of the *Drosophila melanogaster* central nervous system (CNS), also known as neuroblasts, are an excellent model to examine the mechanism driving ACD and has been key in the identification of many of the highly conserved factors involved in this crucial developmental process (Bardin et al., 2004; Zhong and Chia, 2008; Homem and Knoblich, 2012). Neuroblasts divide asymmetrically to generate a self-renewing neuroblast and a ganglion mother cell (GMC), which will further divide to generate neurons or glial cells. The proper establishment of cell polarity is crucial for ensuring the accurate outcome of these divisions and the correct partitioning of cell fate and polarity determinants, which drive the formation of the diverse cell types found in the Drosophila brain. This polarity is regulated by evolutionarily conserved determinants, including Bazooka (Baz, the homolog of mammalian Par-3), Par-6, and atypical PKC (aPKC), known as the Par complex, which localize apically and contribute to the establishment of cell polarity during cell division (Schober et al., 1999; Rolls et al., 2003; Petronczki and Knoblich, 2001). The Par complex also contributes to the correct orientation of the mitotic spindle and division plane through the adaptor protein Inscuteable (Insc) and the apical Gαi / Pins / Mud complex (Kraut et al., 1996; Siller et al., 2006; Nipper et al., 2007). The Scribble (Scrib), Disc Large (Dlg), and Lethal Giant Larvae (Lgl) complex also shares an apical localization and is important for the formation of cell polarity and the correct partitioning of apical / basal cell fate determinants (Albertson and Doe, 2003). The basal cell fate components include the transcription factor Prospero (Pros), Numb, the adaptor protein Miranda (Mira), and the RNA binding protein Staufen (Stau) (Chu-Lagraff et al., 1991; Rhyu et al., 1994; Rolls et al., 2003; Li et al., 1997). It is important to state that these cell fate components are highly conserved among multicellular organisms and the precise nature of the molecular mechanism that is responsible for the proper organization and regulation during cell division of neuroblast is poorly understood.

A potential regulatory mechanism for ACD could be a family of small, non-coding RNAs known as microRNAs (miRNAs). Several miRNAs have recently been identified that have been implicated in a variety of cellular activities including cell differentiation, growth and metabolism (Rottiers and Näär, 2012; Suriya Muthukumaran et al., 2022). miRNAs are known for their ability to silence genes at the post-transcriptional level by binding to the 3’ untranslated region (UTR) of target mRNAs to induce repression and prevent protein translation. miRNAs have been found to be extremely important in a variety of cellular pathways and are generally conserved across all species.

Here, we sought to identify the role of the miRNA, miR-190 in early CNS development. miR-190 have been identified in a host of cellular activities including lifespan, neuronal activity, maintenance, and metabolism, as well as implicated in several diseases including multiple cancers and Alzheimer’s disease (Jia et al., 2016; Ramírez et al., 2022; Yu and Cao, 2019). We performed single cell RNA sequencing (scRNAseq) and comparative transcriptome analysis of miR-190 deficient larval brains and identified several shifts in brain cell populations including a decrease in neurons and increases in neural progenitor cells (NPCs) and glial cells. In addition, in examination of transcriptional profile of miR-190 deficient NPC population, there were several cell fate and polarity determinants that were differentially expressed including Par-6, Lgl, Stau, Pros, aPKC, and Baz. Based on these findings, we shifted our investigation to early CNS development in the Drosophila embryo and found that miR-190 deficient embryos displayed multiple defects in the proper localization of cell fate determinants in mitotic embryonic neuroblast including the mislocalization of the Par complex at the apical cell periphery indicating a lack of establishment of cell polarity. Additionally, many of the basal cell fate determinants were also mislocalized during mitosis, including Pros, Mira, and Stau. Quantitative measurements of putative miR-190 targets in the early embryo displayed significant increases in transcript populations of cell fate and Polarity determinants including Stau, Par-6 and Pros. Notably, we also saw a significant increase in RhoGAP transcript populations indicating a regulatory role in Cdc42 GTPase activation and polarity establishment. Overall, we propose a model that miR-190 targets transcript degradation of cell fate determinants in the cytoplasm to promote activation of Cdc42 leading to the recruitment of the Par complex and the proper establishment of cell polarity.

## Results

### Comparative transcriptome analysis of miR-190 deficient Drosophila larval brains reveal shifts in CNS cell populations and differential expression of several cell fate and polarity determinants

To investigate the role of miR-190 on Drosophila CNS development, we performed scRNA-seq on miR-190^KO^ larval brains to examine differences in cellular composition in the absence of miR-190 (Fig. 1). The miR-190^KO^ line was generated using targeted homologous recombination to remove the hairpin leading to a loss of the mature form of the miRNA (Chen et al., 2014). Homozygous 1st instar miR-190^KO^ larval brains were dissected and dissociated to single cells and sequenced according to previously published approaches (Fig. 1A). We performed quality control (QC) analysis on the data set using the Seurat R package to identify mitochondrial RNA sequences and cells that have unique feature counts (Fig. S1). After quality control and filtering of both datasets, a total of 13,826 cells retained for downstream analysis. Both datasets were integrated and clustered, producing 29 cell clusters which were visualized using Unfold Manifold Approximation and Projection (UMAP) (Fig. 1B, Fig. S2). Clusters were manually annotated based on previously validated marker genes for specific cell types (Fig. 1C, Supplemental Table 1) (Dillon et al., 2022; Brunet Avalos et al., 2019). The 29 clusters were then further grouped into 7 main CNS cell types: neuroblasts, neural progenitor cells (NPCs), undifferentiated neurons (UNs), newborn neurons (NBN), neurons, Kenyon cells (KCs), and glial cells. Two non-neural clusters of hemocytes and muscle cells were also identified. The neuroblast cluster was identified by expression of *deadpan (dpn)*, *miranda (mira)*, and *worniu (wor)* (Dillon et al., 2022). Neural progenitor clusters were identified by *lin-28*, *Notch (N)* and *Tis11* (Komori et al., 2018; Yang et al., 2016; Dillon et al., 2022). Undifferentiated neuron clusters were enriched for *headcase* (*hdc*) and *Thor* while lacking expression of both neural progenitor and mature neuron markers. Newborn neurons were enriched for *Hey* and *E(spl)m6-BFM* (Monastirioti et al., 2010; Michki et al., 2021). Neuron clusters were broadly identified by the expression of neural markers *elav* and *fne*. These clusters were then further specified for mature neural markers including *Gad1*, *VAChT*, *VGlut*, *Vmat*, and *dimm* for the identification of GABAergic neurons, cholingeric neurons, glutamatergic neurons, monoaminergic neurons, and peptidergic neurons, respectively (Dillon et al., 2022). Immature neurons expressed both *elav* and *fne* but lacked robust expression of mature markers (Dilon et al., 2022). Due to the distinct clustering of Kenyon cells apart from the main neuron clusters, we categorized them separately. Kenyon cells were identified by the expression of *prt*, *Dop1R1* and *sNPF* (Brunet Avalos et al., 2019). Glial cells were identified by the expression of *repo* and further specified for astrocyte-like glia, ensheathing glia and surface glia by the expression of *alrm*, *MFS9*, and *Mdr65*, respectively (Xiong et al., 1994; Dillon et al., 2022).

**Figure 1:**
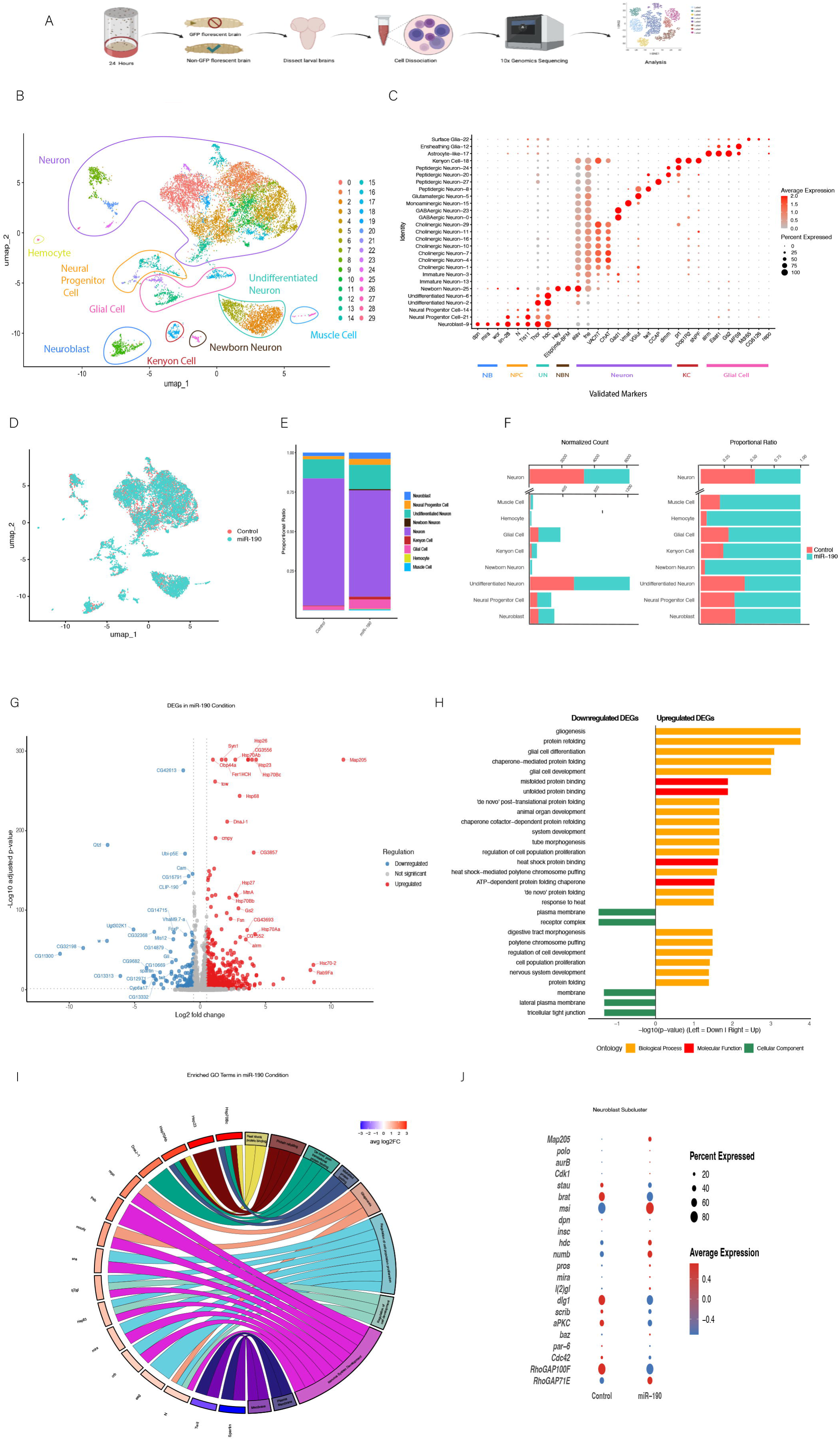
Single-cell RNA-seq workflow and analysis of differentially expressed genes (DEGs) between control and miR-190^KO^ larval brains. (A) Schematic overview of the single-cell RNA-seq sample preparation and data generation process. (B) Uniform Manifold Approximation and Projection (UMAP) of the integrated dataset. Each point represents an individual cell, with cell clusters annotated based on the expression of known marker genes. (C) Dot plot of selected marker genes for each major cell type identified, with dot size representing the percentage of cells expressing each gene and color intensity indicating the average expression level within each cluster. (D) Uniform Manifold Approximation and Projection (UMAP) grouped by condition from the integrated dataset. (E) Bar chart showcasing the proportion of each cell type in control and miR-190^KO^ conditions. (F) Bar chart showing the normalized cell count and proportional ratio of identified cell types across both experimental conditions. (G) Volcano plot of DEGs between control and miR-190^KO^ conditions. Red dots represent genes upregulated in miR-190^KO^ and blue dots represent genes upregulated in control. The y-axis indicates-log10(p-value) and the x-axis shows log2 fold change values. (H) Gene Ontology enrichment analysis of biological processes, molecular function and cellular components for the upregulated DEGs reported in 1G. (I) Chord diagram highlighting the relationship between different DEGs and GO terms. (J) Dot plot depicting the average expression of selected cell fate determinants in the neuroblast cluster between control and miR-190^KO^ conditions. The dot size represents the percentage of cells expressing each gene of interest within the cluster, and the color intensity reflects the average expression level.

To evaluate the effects of miR-190^KO^ on cell fate specification, we analyzed the proportions of different cell types in each condition (Fig. 1D-E, Fig. S3A). Neurons compromised the majority of the control, making up 80.7% of the total cell population. Undifferentiated neurons (12.1%), glial cells (2.3%), neuroblasts (2.3%), neural progenitor cells (1.98%), Kenyon cells (0.42%) and non-neural cells made up the remainder of the control. Cell fate decisions appeared to be altered, as the proportion of neurons in the mIR-190^KO^ condition decreased by 13.2%. This was accompanied by a 3.7% increase in the proportion of glial cells, in addition to increases in undifferentiated cells (3.22%), neuroblasts (2.04%), neural progenitor cells (1.87%), Kenyon cells (1.04%), and newborn neurons (0.5%), respectively. Nearly double the number of cells in the miR-190^KO^ condition relative to the control were captured despite having the same starting number of first instar larval brains. This could indicate that the loss of miR-190 results in increased cell proliferation. To account for differences in cell number, we compared normalized cell counts and proportional ratios of each of the 9 main identified cell type clusters between both conditions (Fig. 1E, Fig. S3B). The number of neuroblasts and neural progenitor cells increased by nearly two-fold in the miR-190^KO^ condition relative to control. A slight increase (1.3-fold) was observed in the normalized cell count of undifferentiated neurons in in the miR-190^KO^ condition relative to control. Notable increases in glial cell and Kenyon cells were also observed, with increases of 2.6-fold and 3.47-fold in the miR-190^KO^ condition relative to control, respectively.

Interestingly, we found that newborn neurons were mainly observed in the miR-190^KO^ condition, representing 96% of the total cell population. As this cluster is enriched for *Notch* responsive genes (*Hey*, *E(spl)m6-BFM*, *E(spl)m8-HLH*) and genes involved in asymmetric cell division (*pros*, *polo*, *CDK1*), this cluster represents Type A neurons resulting from asymmetric cell divisions of ganglion mother cells (GMC) (Monastirioti et al., 2010). This data suggests that miR-190 may be involved in maintaining the proper balance of cell populations during CNS development, potentially through the regulation of asymmetric cell division.

Based on our findings involving the shifts in CNS cell populations, we sought to examine more specific changes at the transcriptional level. We examined the differential gene expression between the miR-190^KO^ transcriptome and the control transcriptome (Fig. 1G). A total of 633 genes were identified as differentially expressed genes (DEGs), with an adjusted p-value < 0.05 and absolute log2 fold change (log2FC) > 0.58 (Supplemental Table 2). Of these, 518 were upregulated and 115 were downregulated.

Notably, one of the top DEGs, microtubule associated protein 205 (Map205), was significantly upregulated in the miR-190^KO^ population. Map205 has been shown to interact and inhibit Polo kinase providing a stable reservoir of Polo for entry into mitosis (Archambault et al., 2008). In addition, the gene syntrophil1 (syn1) also was highly expressed in miR-190^KO^ condition. Syn1 is part of a complex of proteins known as the dystrophin glycoprotein complex (DGC) which includes dystroglycans and sarcoglycans that have been found to be necessary for G-protein signaling pathways (Xiong et al., 2009; Zhou et al., 2006). Interestingly, DGC has also been implicated in the regulation of miRNA levels in the cell (Marrone et al., 2012). Gene Ontology (GO) analysis using GProfiler revealed that upregulated DEGs were significantly enriched for biological process (BP) terms associated with glial cell development (gliogenesis, glial cell differentiation, glial cell development), BP and molecular function (MF) terms associated with protein folding and binding (chaperone-mediated protein folding, protein refolding, and heat shock protein binding), and BP terms associated with the regulation of cell proliferation and development (regulation of cell development, cell population proliferation, nervous system development) (Fig. 1H, Supplemental Table 3). Fig. 1I, Fig. S4 illustrates the distribution of significantly upregulated DEGs of interest across various Gene Ontology (GO) terms. Heatshock proteins including Hsp70Bc, Hsp70Ab, Hsp23 and DnaJ-1 were found across multiple GO terms associated with protein binding. This indicates the presence of cellular stress, which is in line with the nature of the miR-190^KO^ allele, as it displays a lethality in 3rd instar larvae (Chen et al., 2014). Interestingly, Hsp83 was shown to be classified with GO terms associated with regulation of cell proliferation and development. *Repo*, *N* and moody were found in both gliogenisis and nervous system development terms. Notably, *esg*, *sna*, *l(2)gl*, *crumbs (crb)*, and *miranda (mira)*, genes involved in the regulation of cell polarity and neuroblast asymmetric cell division, were found in the regulation of cell population proliferation, nervous system development, and regulation of cell development terms. Increased expression of these genes in the miR-190^KO^ condition relative to the control indicate that explain the changes in cell fate seen between both experimental conditions. Downregulated genes were found to only be significantly enriched for cellular component terms associated with the plasma membrane (membrane, lateral plasma membrane, plasma membrane), receptor complex and tight junctions (tricellular tight junction) (Fig. 1H, Supplemental Table 3). Target of Wit (twit) and Spartin, two genes involved in regulating the BMP pathway, were found in the membrane and plasma membrane terms (Fig. 1I). Significant downregulation of these genes may explain cell fate changes seen in the miR-190^KO^ condition, as Spartin was found to regulate neuronal survival through its role in modulating microtubule stability via the BMP signaling pathway (Nahm et al., 2013).

To further investigate the impact of miR-190^KO^ on the expression of other cell fate and polarity determinants, we subclustered the neuroblast cluster and compared the expression profiles between both conditions (Fig. 1J). Our analysis revealed altered gene expression profiles in the miR-190^KO^ compared to control conditions. Notably, genes such as *mira*, *Musashi (msi)*, and *prospero (pros)* were more highly expressed in miR-190^KO^ conditions (Fig. 1I). Conversely, genes such as *dlg1*, *Stau*, *aPKC*, and brat were more highly expressed in control conditions. Overall, our comparative transcriptome analysis demonstrated that miR-190^KO^ larval brains display a shift in CNS cell populations, with notable decreases in neurons and increases in glial cells, neuroblasts, Kenyon cells, undifferentiated cells, NPCs and newborn neurons. In addition, significant changes in the expression of cell fate and polarity determinants indicate a role for miR-190 in regulation of cell fate determinants and the generation of CNS cell diversity.

### qPCR of Drosophila embryos deficient for miR-190 demonstrates elevated levels of several cell and polarity fate determinants

Based on our scRNA-seq data, we hypothesized that miR-190 is involved in cell fate determination, potentially acting early in development, as suggested by the differential expression of several cell fate and polarity determinants (Fig. 1I). In support of this hypothesis, we also examined the populations of neuroblasts and GMC-fated cells in first instar miR-190^KO^ larval brains using immunofluorescence analysis. We found a significant increase in GMC-fated cells and a corresponding decrease in neuroblasts (Fig. S4).

To investigate the potential role of miR-190 in embryonic CNS development, we identified its putative targets using the predictive software TargetScanFly 7.2 (Agarwal et al., 2015). TargetScanFly predicts regulatory targets of miRNAs by identifying mRNAs with conserved complementary sites in their 3′ UTRs. miR-190 exists in two isoforms miR-190-5p and miR-190-3p, with TargetScanFly identifying 331 and 498 potential targets, respectively. We found that over a third of the predicted targets are involved in neurogenesis and neural function. Notable neurodevelopment-related targets are listed in Supplementary Table 4. We found that miR-190 displays conserved 8-mer matches to the transcription factor *pros*, the tumor suppressor *scribble* (*scrib*), the cell polarity factor *Par-6*, the neural RNA-binding protein *Musashi* (*msi*), and several *RhoGAPs*. Based on these predicted targets, we further investigated the impact of miR-190^KO^ on transcript levels of known cell fate and polarity determinants in the developing embryo. We performed quantitative reverse transcriptase PCR (qRT-PCR) on miR-190^KO^ embryos to assess expression levels of conserved molecular determinants (Fig. 2). Embryos at developmental stages 9–12 were used for this analysis.

**Figure 2.**
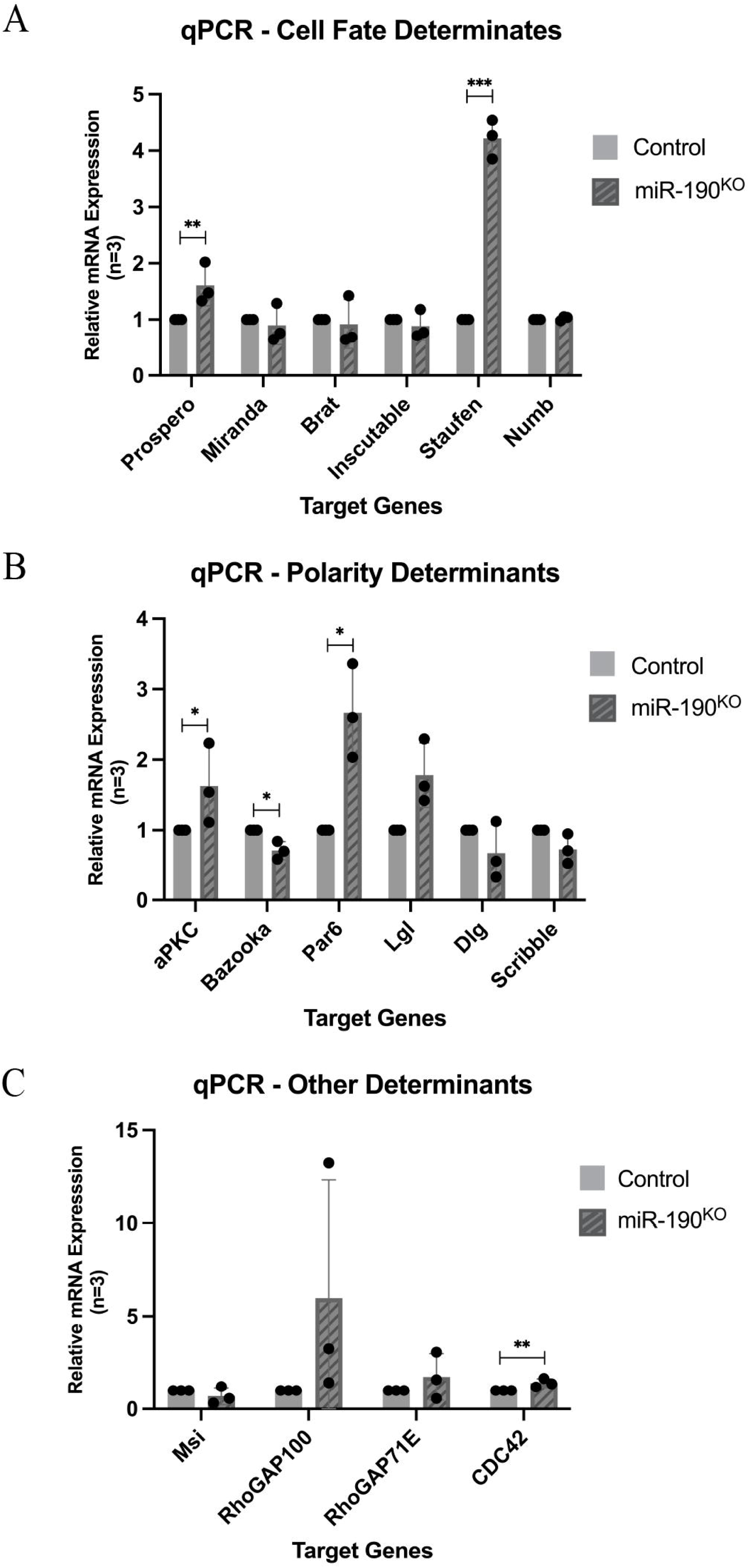
Several cell fate, cell polarity, and molecular determinants are upregulated in miR-190-deficient Drosophila embryos. Relative mRNA expression patterns of established cell fate, cell polarity, and molecular determinants between control and miR-190^KO^ from stage 10-12 *Drosophila* embryos. (A) cell fate determinants prospero was upregulated almost two-fold, while Staufen was upregulated almost five-fold. (B) cell polarity determinants Lgl were upregulated almost two-fold, while Staufen was n=3 experimental repeats; asterisks indicate p≤ 0.05. Bars represent the means and standard errors of three biological replicates. Error bars: standard deviation; *p≤0.05; **p≤ 0.005; ****p≤ 0.0005.

We organized the conserved targets into three functional categories: cell fate determinants (Fig. 2A), cell polarity determinants (Fig. 2B), and other molecular determinants involved in polarity and fate, but not acting directly (Fig. 2C). Among the cell fate determinants, *pros*, *lgl*, and *Stau* showed significantly altered mRNA expression levels (Fig. 2A). *Pros* exhibited nearly a twofold increase in expression in miR-190^KO^ embryos (*p* = 0.004), while *Stau* expression increased fivefold (*p* = 0.0007) compared to controls. Similarly, *lgl* mRNA levels nearly doubled in miR-190^KO^ embryos (*p* < 0.0415), whereas *dlg* and *scrib* showed slightly reduced expression. Analysis of cell polarity determinants (Fig. 2B) revealed higher mRNA levels of *aPKC* (*p* = 0.04) and *Par-6* (*p* = 0.0202) in miR-190^KO^ embryos, while *baz* expression was slightly reduced (*p* = 0.0478). For other molecular determinants (Fig. 2C), *RhoGAP100* (*p* = 0.2445) and *Msi* (*p* = 0.3655) were significantly upregulated, while *Deadpan*, *RhoGAP71E*, and *Cdc42* expression levels remained comparable to controls. Several of our qPCR results are consistent with the predicted targets of miR-190, including *pros*, *RhoGAP100*, *Par-6*, and *Msi* (Supplementary Table 4). However, *Stau* and the cell polarity factor *Lgl* were not predicted to directly bind miR-190, suggesting they may be regulated indirectly via other miR-190 targets. We conclude that miR-190 regulates both cell polarity and fate selection by directly targeting the transcripts of *Par-6*, *RhoGAP100*, and *pros*, and by indirectly influencing *Stau* and *Lgl* levels. Taken together, our data suggest that miR-190 is involved in establishing proper cell polarity and maintaining the balance of cell fate determinants during development.

### The basal cell fate determinants Pros and Mira are no longer restricted to the basal cortex in miR-190^KO^ mitotic embryonic NBs

Based on our scRNA seq data, we found that there were significant shifts in larval brain cell populations, indicating a possible role for miR-190 in development and the generation of cell type specification. To examine this, we focused on CNS development in late-stage Drosophila embryos (stage 12). We were interested in investigating whether cell fate determinants were being partitioned correctly during neuroblast cell division. We fixed and immunostained miR-190^KO^ late-stage embryos for the basal cell fate determinant Pros and the adapter protein, Mira and examined their distribution during mitosis (Fig. 3). In control embryos, upon entry into mitosis, Pros localizes at the cortex and then accumulates at the basal side of the cell during metaphase forming a crescent (Fig. 3A, top panels). Similar to Pros, Mira displayed a basal crescent formation during metaphase. This localization at the basal side of the cell allows Pros and Mira to be inherited into one of the daughter cells during cell division driving differentiation. In miR-190^KO^ embryos, Pros did not form a basal crescent at metaphase but rather was found localized along the cell periphery during anaphase (Fig. 3A, bottom panels). Additionally, Mira did not localize as a basal crescent, rather it displayed a similar localization at Pros along the cell periphery. To better understand the penetrance of both the Pros and Mira mislocalization phenotypes, we performed counts of cells, in both control and miR-190^KO^ embryos, that displayed a basal Pros and Mira crescent during mitosis (Fig. 3B). An average of ∼6 - 10 embryos were counted for each experimental repeat (N=3). In control embryos, ∼85 embryonic neuroblasts displayed a basal Pros crescent during mitosis, however in miR-190^KO^ only 12 mitotic cells showed a basal Pros crescent at anaphase (Fig. 3B). Additionally, 35 embryonic neuroblasts displayed Pros uniform localization along the cortex, no longer restricted to the basal end of the cell during anaphase. For control counts of Mira localization 87 neuroblasts displayed a proper basal crescent formation of Mira during mitosis, in contrast with 37 neuroblasts displaying a uniform localization along the cortex of Mira lacking a basal localization during mitosis in miR-190^KO^ embryos (Fig. 3C). Here, we conclude that both the transcription factor Pros, and its adapter protein Mira are mislocalized during mitosis in miR-190^KO^ embryos.

**Figure 3.**
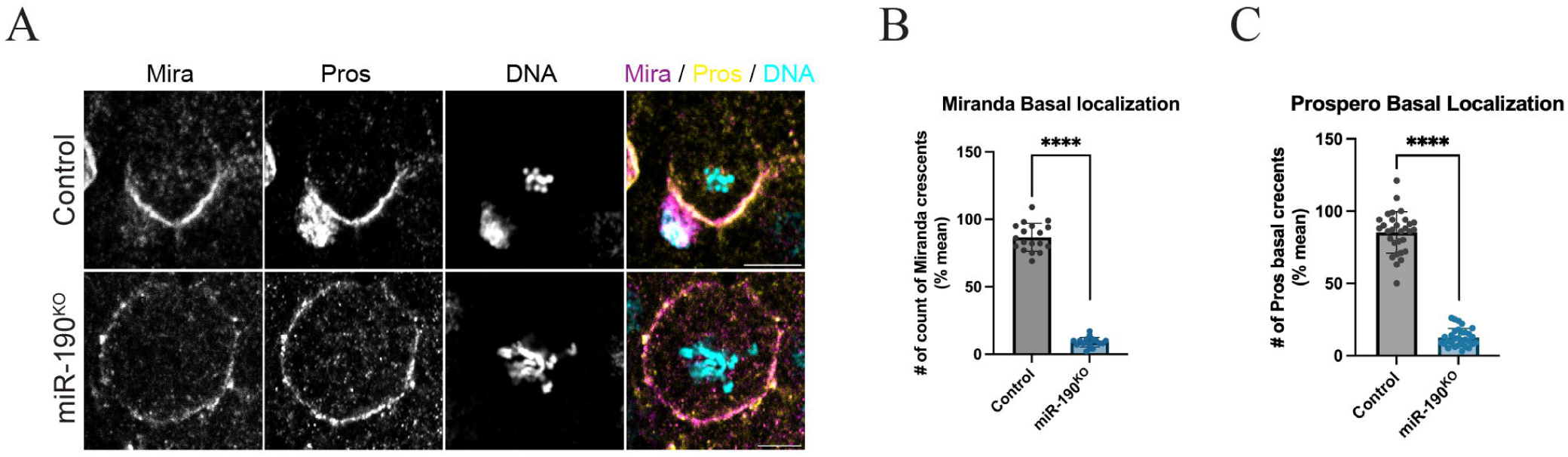
Prospero and Miranda display cortical localization during mitosis in miR-190 deficient embryos. (A) Control and miR-190^KO^ deficient embryos were stained during embryonic stages 10-12 for the basal cell fate determinants, Pros (yellow), Mira (magenta) and DNA (cyan). During metaphase, control embryos display a crescent localization at the basal cortex, however in miR-190 embryos at the same stage during mitosis, both Mira and Pros show a cortical localization at the cortex, and a lack of crescent formation (bottom panels), Scale bars: ∼5μm. (B) Counts of mitotic NB cells in control and miR-190 embryos display a significant lack of crescent formation of Mira during mitosis. (C) Counts of Pros crescent formation in mitotic NB display a significant lack of proper crescent formation at metaphase in miR-190^KO^ embryos. n=3 experimental repeats, 6-8 embryos per experiment. Mean±s.e.m., **** P≤0.0001.

### The cell polarity Par complex (aPKC, Baz, and Par-6) are mislocalized in miR-190^KO^ embryonic mitotic neuroblasts

Because of the mislocalization of the basal cell fate determinants, Pros and Mira, we sought to investigate whether cell polarity is established in miR-190^KO^ embryonic neuroblasts. We examined the localization of the Par complex during mitosis miR-190^KO^ embryos (Fig. 4). The highly conserved Par complex consists of the PDZ domain proteins Baz / Par-3, Par-6, and the kinase aPKC and are integral for the establishment of cell polarity in a variety of tissues and cell types (Wodarz, 2002; Vorhagen and Niessen, 2014). Furthermore, establishment of cell polarity is critical for the correct partitioning of cell fate determinants during asymmetric cell divisions. We fixed and immunostained control and miR-190^KO^ embryos during stages 9-12 and examined the correct localization of the Par complex proteins during mitosis in embryonic neuroblasts. In our controls, Baz forms a crescent localization at the apical domain and Pros displays a crescent localization at the basal end of the cell during mitosis (Fig. 4A, top panels). However, in miR-190^KO^ embryos, we see mislocalization of Baz during mitosis where it is no longer at the apical periphery, rather it appears to be largely cytoplasmic and punctate and co-localized with Pros rather than displaying an opposite apical / basal localization (Fig. 4A, bottom panels). We also immunostained control and miR-190^KO^ stage 9-12 embryos for aPKC and found that while in control mitotic embryonic neuroblasts, aPKC forms an apical crescent, in miR-190^KO^ embryonic neuroblasts, aPKC appeared diffuse and largely spread along the cell periphery (Fig. 4B). Lastly, we also examined the localization of PDZ domain protein Par-6 in control and miR-190 deficient embryos (Fig. 4C). Similar to the other Par complex proteins, Par-6 also displayed an apical crescent formation upon entry into mitosis (Fig. 4C, top panels). However, in miR-190^KO^ embryonic mitotic NBs, Par-6 appeared punctate and largely diffused along the periphery of the cell (Fig. 4C, bottom panels). We also performed counts of mitotic apical crescent formation of Baz, aPKC, and Par-6 in control and miR-190^KO^ embryonic NBs (Fig. 4D, E, and F). In control embryos, 69 embryonic NB showed an apical Baz crescent, while in miR-190^KO^ embryos, 11 embryonic NB displayed an apical crescent (P≤0.005) (Fig. 4D). Interestingly, 43 embryonic NB showed a cytoplasmic and punctate distribution of Baz during metaphase. Counts involving aPKC mitotic crescent formation showed that in control embryos, the large majority of NBs exhibited proper apical localization of aPKC, while in miR-190^KO^ embryos, this number dropped significantly (Fig. 4E; P≤0.0005). This was also true for Par-6, with a significant lack of Par-6 mitotic apical crescents formation in miR-190^KO^ embryonic NBs (Fig. 4F; P≤0.0005). Here we conclude that miR-190^KO^ is necessary for the correct localization of the Par complex during mitosis, indicating that cell polarity is not established, and thus other basal cell fate determinants that rely on cell polarity may be affected.

**Figure 4.**
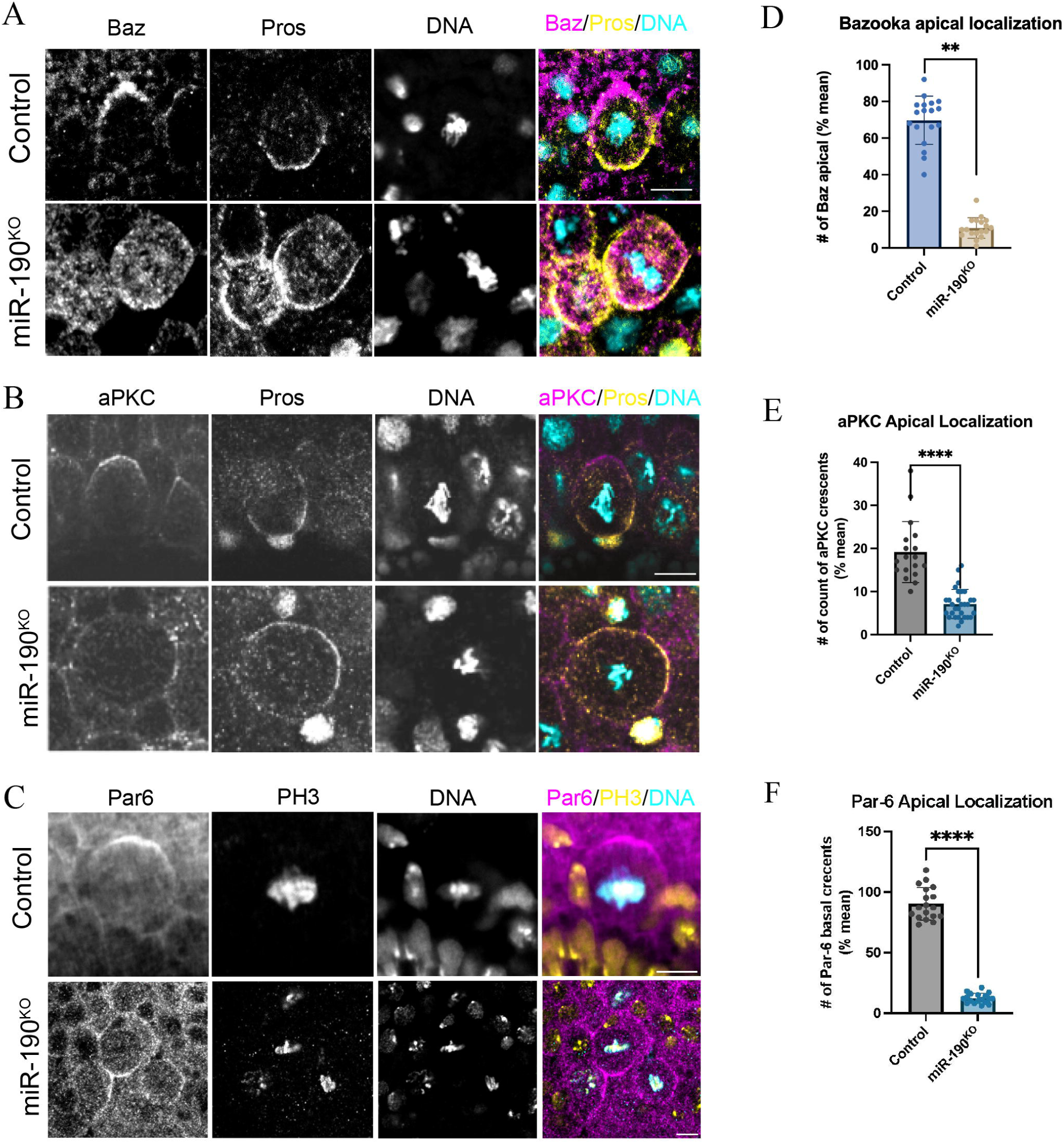
The PAR complex is mislocalized in miR-190 deficient embryonic NB. (A) Control and miR-190^KO^ embryos were fixed and immunostained for Baz (magenta), Pros (yellow), and DNA (cyan). At metaphase, Baz displays an apical crescent localization, while Pros displays a basal crescent localization (top panels). However, in MiR-190^KO^ embryonic NBs at metaphase, Baz is mislocalized displaying a punctate distribution throughout the cell, while Pros displays a cortical localization (bottom panels).Scale bar: ∼5μm. (B) Control and miR-190^KO^ embryonic NBs were stained for aPKC (magenta), Pros (yellow), and DNA (cyan) during metaphase. At metaphase, aPKC displays an apical crescent localization, while Pros displays a basal crescent localization (top panels). In miR-190^KO^ embryonic NBs, aPKC displays a diffuse cortical localization during mitosis and Pros shows a cortical localization (bottom panels). Scale bar: ∼5 μm. (C) Control and miR-190^KO^ embryonic NBs were immunostained for Par6 (magenta), PhosphoH3 (yellow), and DNA (cyan) during metaphase. Par6 shows an apical cortical localization during metaphase, while in miR-190^KO^ embryonic NBs there is a diffuse cortical staining. Scale bars: ∼5μm. (D) miR190^KO^ embryonic NB displays a decrease in Baz mitotic crescent formation. The bar graph represents mean counts of Baz crescents in control (blue) and miR-190^KO^ (brown) stage 10-12 embryos during mitosis. There is a significant decrease in mitotic apical crescent formation of Baz in miR-190^KO^ embryonic NBs. (E) miR190^KO^ embryonic NB displays a decrease in aPKC mitotic crescent formation. The bar graph represents mean counts of aPKC crescents in control (blue) and miR-190^KO^ (brown) stage 10-12 embryos during mitosis. There is a significant decrease in mitotic apical crescent formation of aPKC in miR-190^KO^ embryonic NBs. (F) miR190^KO^ embryonic NB displays a decrease in Par6 mitotic crescent formation. The bar graph represents mean counts of Par6 crescents in control (blue) and miR-190^KO^ (brown) stage 10-12 embryos during mitosis. There is a significant decrease in mitotic apical crescent formation of Par6 in miR-190^KO^ embryonic NBs. n=3 experimental repeats, 6-8 embryos per experiment. Mean ± s.e.m., **P≤0.005, ***P≤0.0005 ****P≤0.00005.

### The cell polarityprotein Scribble displays a diffuse cortical localization in miR-190^KO^ embryonic neuroblast

Scribble (Scrib), with Dlg and Lgl are a second protein complex known to regulate cell polarity in epithelia and neuroblast (Bilder and Perrimon, 2000; Albertson and Doe, 2003; Peng et al., 2000). To determine the effect of miR-190 on the localization of Scrib, we performed immunofluorescence staining of dividing neuroblast in miR-190^KO^ embryos. In control embryos, Scrib displayed an apical-basolateral cortical localization during metaphase and is diffuse at the basal cortex (Fig. 5A, top panels). However, in miR-190^KO^ embryos, Scrib displayed a diffuse uniform distribution around the cell cortex throughout mitosis (Fig. 5A, bottom panels). Quantitative analysis showed that while control embryos exhibited correct Scrib localization, the number of embryos with proper Scrib localization dropped significantly in miR-190^KO^ embryos (Fig. 5B; P≤0.005). These results are significant because Scrib is a key scaffolding protein involved in establishing and maintaining apical-basal polarity within neuroblasts. The mislocalization of Scrib in miR-190^KO^ embryos suggests that miR-190 is necessary for the proper localization of Scrib, and by extension, the establishment of polarity.

**Fig. 5.**
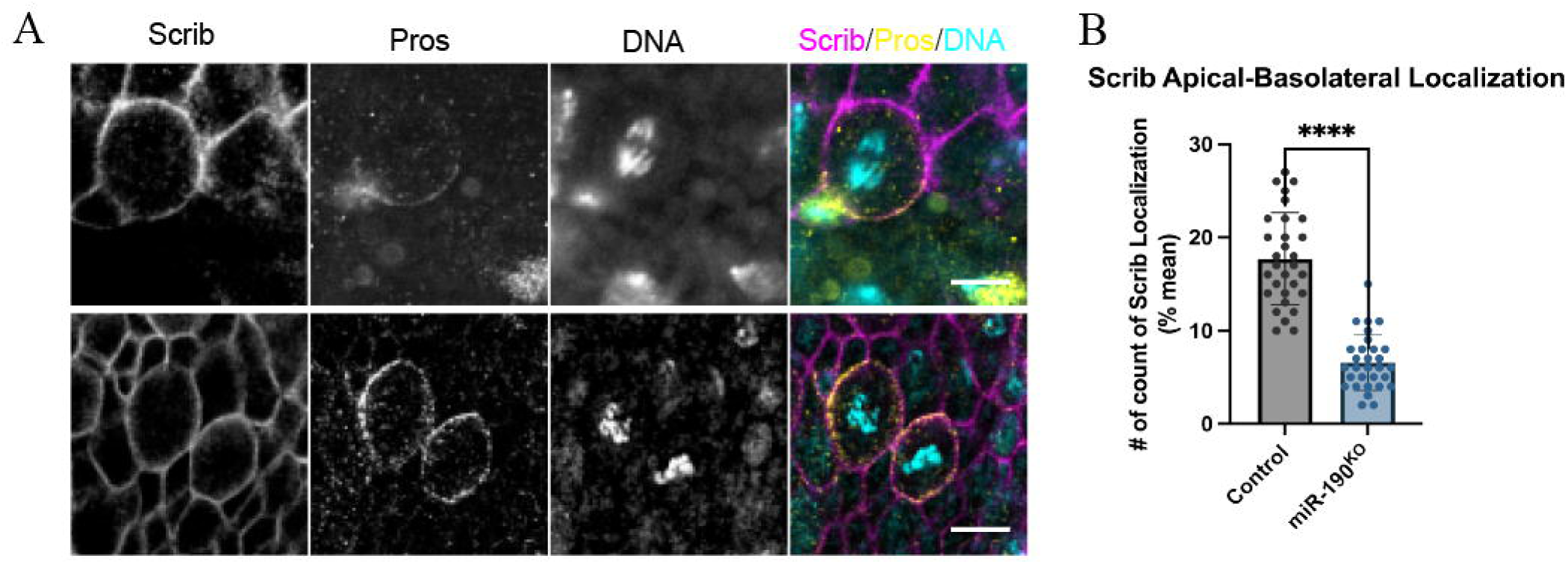
Scrib displays a cortical localization along the cell periphery in miR-190^KO^ embryos. (A) control and miR-190^KO^ stage 12 embryos were fixed and immunostained for Scrib (magenta), Pros (yellow) and DNA (blue). Upon entry into mitosis, Scrib displays an apical-basolateral localization (top panels), however in miR-190^KO^ embryos, Scrib shows a uniform localization along the cortical membrane and does not display an accumulation along the apical-basolateral cortex (bottom panels). (B) Counts of Scrib apical-basolateral crest in mitotic embryonic NB in control and miR-190^KO^ embryos. There is a significant loss of apical-basolateral Scrib crescents in mitotic NBs in miR-190^KO^ embryos. n=3 experimental repeats, 6-8 embryos per experiment. Mean ± s.e.m., **P≤0.005.

### The RNA-binding protein Staufen mislocalizes to the mitotic spindle in miR-190^KO^ neuroblasts

Our findings displayed that the scaffold protein, Mira and its cargo, Pros are mislocalized in miR-190^KO^ embryonic neuroblasts (Fig. 3). We sought to investigate the effects of other Mira-related cargoes in miR-190 deficient embryos. We immunostained control and miR-190KO stage 12 embryos for the double stranded RNA-binding protein Stau,-tubulin, and the mitotic marker Phospho-H3 (Fig. 6). Previous studies have shown that Stau and Mira interact, and that Mira is necessary for Stau basal crescent localization (Matsuzaki et al., 1998; Jia et al., 2015). In control embryos, we found that Stau showed a basal crescent localization in mitotic neuroblasts (Fig. 6A, top panels). Surprisingly, in miR-190^KO^ mitotic neuroblasts, Stau no longer formed basal crescents but rather aberrantly localized along the mitotic spindle (Fig. 6A, bottom panels). We also noticed that there were defects in mitotic spindle rotation along the apical / basal axis in miR-190^KO^ embryonic neuroblasts (Fig. S5). We performed counts of Stau basal crescent formation in control and miR-190^KO^ embryos and found that there was a significant lack of Stau crescent formation in miR-190^KO^ embryonic NBs (Fig. 6B; P≤0.0005). Here we conclude that Stau is mislocalized in miR-190^KO^ neuroblasts, displaying an accumulation along the mitotic spindle, due to lack of cell polarity.

**Fig. 6.**
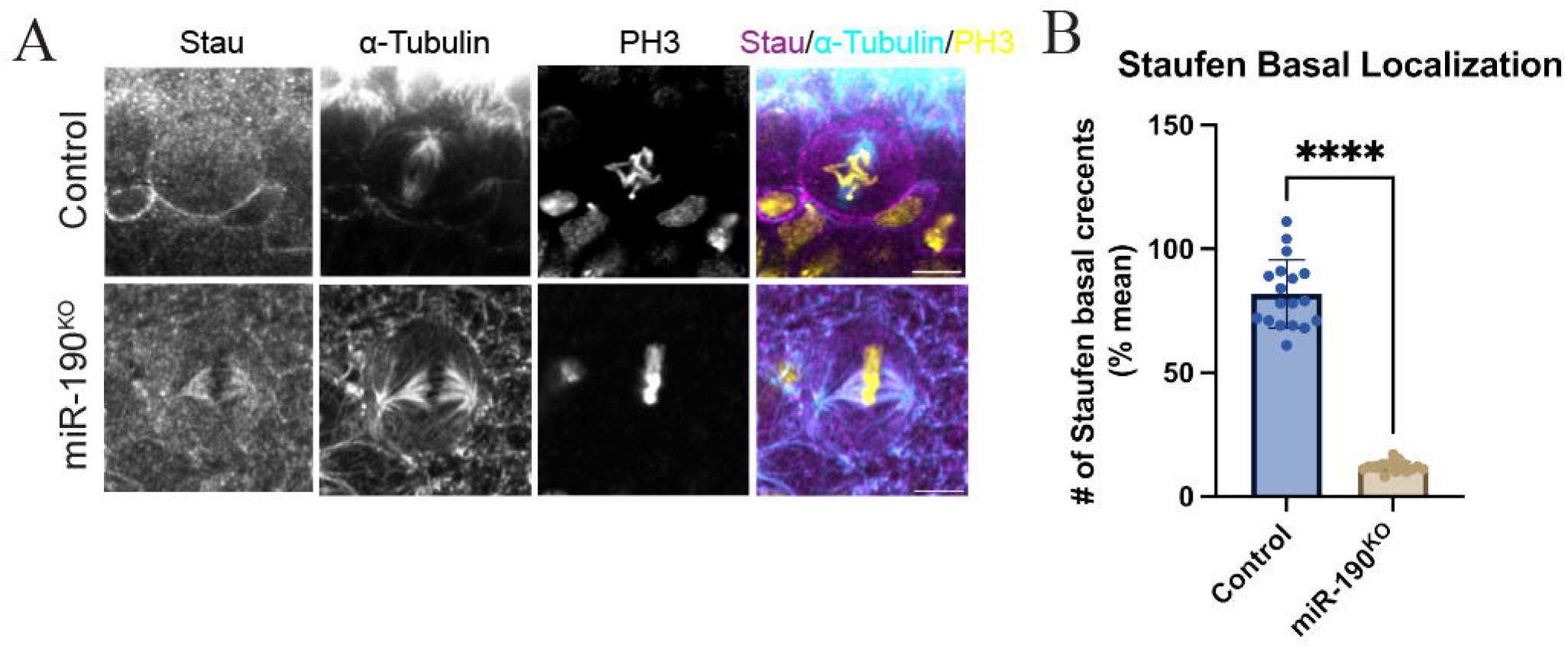
Staufen displays a localization along the mitotic spindle in miR-190 deficient embryos. (A) Control and miR-190^KO^ embryos were fixed and immunostained for the cell fate determinant Stau (magenta),-tubulin (cyan), and phospho-H3 (yellow) during mitosis in embryonic NBs. In control embryos, stau displays a basal crescent localization at metaphase (top panels), however in miR-190^KO^ embryonic NBs Stau localizes along the mitotic spindle during metaphase (bottom panels) scale bar: ∼5μm. (B) Counts of Stau mitotic crescent formation in control and miR-190^KO^ embryonic NBs. There is a significant decrease in Stau crescent formation in mitotic NB in miR-190^KO^ embryos. n=3 experimental repeats, 6-8 embryos per experiment. Mean ± s.e.m., ****P≤0.0005.

## Discussion

Here, we report that miR-190 is a key regulator of cell polarity in dividing neuroblast by directly targeting Pros, Scrib, RhoGAP transcripts during cell fate selection (Fig. 7A). In addition, miR-190 is required for proper establishment of cell polarity, as the Par complex (Baz, Par-6, aPKC) are mislocalized in the absence of miR-190. In turn, several of the cell fate determinants are also mislocalized due to a lack of cell polarity in miR-190^KO^ embryonic NBs. Here, we propose a model that miR-190 targets several factors that both affect the establishment of cell polarity and displays defects in the proper localization of several key cell fate determinants including Baz, Par-6, Pros, and Stau (Fig. 7B). Additionally, due to the lack of cell polarity, there are defects seen in proper apical-basal mitotic spindle alignment during metaphase (Fig. 7B, Fig. S5).

**Figure 7.**
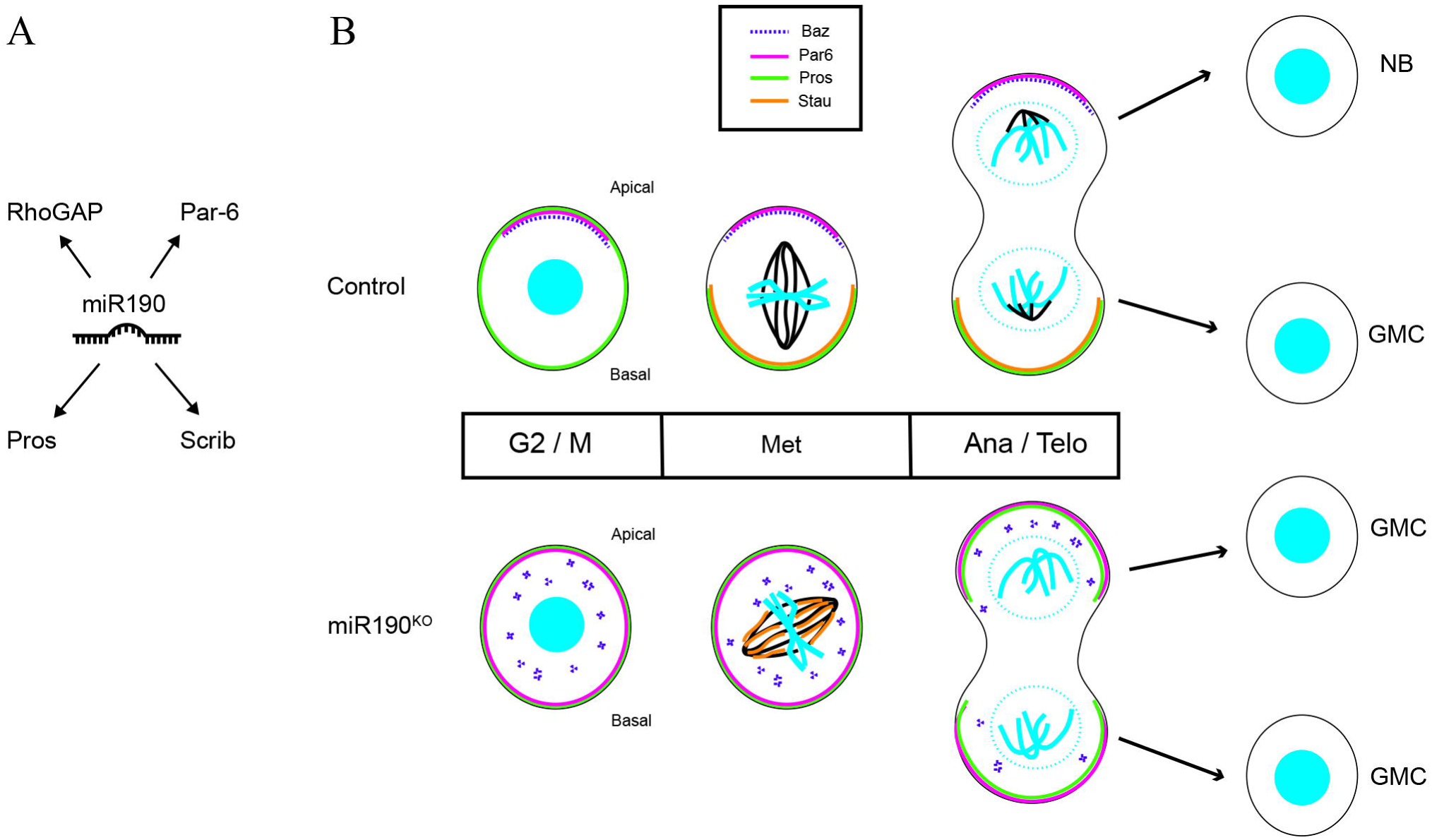
miR-190 targets key cell fate determinants for the correct establishment of cell polarity and cell fate selection. (A) Based on *in silico* predictions, scRNAseq, and qPCR approaches, we propose that miR-190 targets several factors prior to entry into mitosis, allowing for the proper establishment of cell polarity including Scrib, RhoGAP, Pros, and Hdc. (B) In control cells, miR-190 activity allows for the proper establishment of cell polarity, which includes proper apical localization of the Par Complex (Baz; dotted purple line, Par-6; magenta line), which allows for the proper localization of Pros (green line) and Stau (orange line) at the basal cortex. Correct alignment of the mitotic spindle occurs at metaphase, and subsequently the daughter cells adopt the proper cell fate of a self-renewing NB and a differentiated GMC. In contrast, in miR-190^KO^ cells, early in mitosis Baz is mislocalized in the cytoplasm, while Par-6 and Pros localize along the cortex and does not form a crescent apically or basally. Additionally, Stau does not form a basal crescent during mitosis, rather is mislocalized along the mitotic spindle. Upon exit from mitosis, based on the incorrect partitioning of Pros and the lack of NB in larval brain cells, miR-190^KO^ both daughter cells adopt a differentiated GMC state.

### scRNA sequencing of a miR-190^KO^ Drosophila larval brain transcriptome displayed shifts in cell populations and a differential expression of several cell fate and polarity determinants

Based on the shifts in neural cell populations (Fig. 1D, E) and the lack of neuroblast in the miR-190^KO^ larval brains (Fig. S4), we hypothesized that miR-190 played an important role in neural development and sought to understand how larval brain development is affected in miR-190^KO^ larvae. It is important to note that homozygous miR-190^KO^ lines display lethality at late instar development (Brunet Avalos et al., 2019; Dillon et al., 2022). We performed scRNA sequencing of 1st instar larval brains and sought to identify major shifts in transcript populations and cell fate during development. We found that there were increases in glial, neural progenitor cells (NPCs) and undifferentiated cell populations and a decrease in neuron cell populations in miR-190^KO^ larval brains (Fig. 1). This finding is in line with a previous study that created small RNA-Seq libraries for neuroblast, neurons and glia and identified miR-190, as being upregulated in these populations (Menzel et al., 2019). Interestingly, miR-190 has also been identified in several other studies involving life span, regulation of hypoxia and immune response (Li et al., 2014; De Lella Ezcurra et al., 2016; Jiang et al., 2018). In a recent study identifying neurotransmitter specification involving development of specific neuron populations, identified several non-coding RNAs including miR-190 (Estacio-Gómez et al., 2020). Based on our findings and other studies involving neural development and behavior, we believe that miR-190 plays an important role in central nervous system development (CNS). It is important to point out that many of the aforementioned studies in Drosophila involved screens in which miR-190 was identified, along with other microRNAs suggesting that miR-190, while playing an important role, may work in conjunction with other microRNAs to guide neural development.

In examination of expression of cell fate transcripts of the NB cluster, several targets emerged as differentially expressed in miR-190^KO^ larvae. In particular, RhoGAP71E and RhoGAP100 showed a strong increase in expression in miR-190^KO^ NB populations. Taken together with the binding prediction of miR-190 interacting with RhoGAP71E and RhoGAP100, this allowed us to hypothesize that miR-190 is directly regulating RhoGAP activity shifting the population of Cdc42 to an active GTP state (Fig. 7). This is in line with studies that have shown that Cdc42 recruits Par6-aPKC to the apical cell periphery (Joberty et al., 2000; Kay and Hunter, 2001; Lin et al., 2000). Higher RhoGAP levels, resulting in lower Cdc42 activity may ultimately cause the defects in Par6-Baz-aPKC localization seen in embryonic neuroblasts. Another interesting target that was highly expressed in miR-190^KO^ neuroblasts is Hdc, the Drosophila ortholog of the human tumor suppressor HECA (Makino et al., 2001). Hdc has been linked to various developmental processes including imaginal disc morphogenesis, maintenance of the niche in the testis and intestinal stem cells, and as a novel tumor marker for colorectal cancer (Resende, 2013; Resende et al., 2013; Chien et al., 2006). A recent study has also linked Hdc to insulin / mTOR signaling and regulates the timing of neuronal differentiation (Avet-Rochex et al., 2014). It would be interesting to examine if miR-190 targets Hdc in establishing proper timing in CNS development. The neural RNA binding protein Msi is also a very intriguing target, as it is required for asymmetric cell division of the sensory organ precursor cells (SOP) in the adult Drosophila sensory organ (Nakamura et al., 1994). During sensory organ development, the SOP asymmetrically divides to give rise to a non-neuronal precursor cell and a neuronal precursor cells. In msi deficient flies, asymmetric division fails and two non-neuronal precursor cells are produced. Msi has been proposed to regulate this planar cell polarity through post-translational control of transcription and interaction with miR-190 could indicate a general role in regulation of asymmetric cell division.

### miR-190 is a key regulator in establishment of cell polarity in differentiating neuroblast populations

Our scRNAseq transcriptome data suggest that miR-190 may play a role in cell differentiation, therefore we investigated early type I neuroblast division during embryogenesis. Using Targetscan, we identified that miR-190 displayed a strong complement to the multi-PDZ domain protein Scrib and the transcription factor Pros (Supplemental table 3). We investigated transcript levels of known cell fate determinants involved in cell polarity, cell differentiation and stemness in miR-190^KO^ embryos. We identified Staufen, Lgl, Pros, aPKC, and Scribble displaying significant changes in transcript level in the absence of miR-190 (Fig. 5). In addition, we see a mislocalization of aPKC and Pros, as well as the coiled coil protein Mira during cell division of neuroblast in miR-190^KO^ embryos (Figs. 6, 7). Specifically, we see a localization along the cortex, no longer confined to the mitotic crescents. The mislocalization of aPKC is likely due to the lack of cell polarity establishment in dividing neuroblasts, as there is also a mislocalization of Bazooka / Par-3, including a loss of apical crescents at the cortex and a punctate distribution in the cytoplasm (Fig. 4A).

Prior studies showed that *in vitro*, the Par components bind to each other and *in vivo*, they are interdependent for their proper localization in neuroblasts (Lin et al., 2000; Wodarz et al., 2000). A loss of cell polarity during neuroblast division will greatly affect both the proper partitioning and inheritance of apical and basal cell fate determinants. Proper apical / basal localization of these determinants allows for the activity of these proteins to be limited to a specific region of the cell. This is especially true of aPKC, as its activity at the apical region of the cell allows for the correct establishment of many basolateral components. An earlier study of aPKC zygotic null mutants showed that in embryonic neuroblasts, in the absence of aPKC, there is a lack of apical localization of Lgl and a failure to exclude Mira from the apical cortex (Atwood and Prehoda, 2009). These are similar phenotypes that we observe in miR-190^KO^ embryonic neuroblasts, indicating that aPKC cannot restrict its activity to the correct region of the cell.

### miR-190 as a regulator of neurogenesis and cancer progression

Our data indicates that miR-190 plays an important role in the establishment of cell polarity and cell fate in dividing neuroblast during development. This is evident by the mislocation of key polarity complexes including the Par complex and the Scrib/Lgl complex and the targeting of the transcription factor, Pros. Drosophila miR-190 has been identified in several screens related to hypoxia response, neurogenesis, and neural maintenance pathways (De Lella Ezcurra et al., 2016; Fernandes and Varghese, 2022). Interestingly, the human homolog, hsa-miR-190 has also been implicated in neuronal differentiation and a variety of cancers (Jia et al., 2016; Liu et al., 2024; Yu and Cao, 2019). While there may be some differences in sequence and function between species, the conservation suggests that these miRNAs might play similar roles in regulating gene expression in neuronal health and development. Our study outlines a fundamental role for miR-190 in cell polarity that could shed light on the mechanism involving its role in cancer progression.

Defects in the Scrib/Lgl/Dlg complex activity have been directly linked to cancer development, as it modulates Ras-mitogen activated protein kinase (MAPK) signaling activity (Dow et al., 2008). The human homologs of the Scrib/Lgl/Dlg complex have all been shown to be mislocalized in a variety of cancers and have been defined as neoplastic tumor suppressors (Gardiol et al., 2006; Feigin et al., 2014; Pearson et al., 2011). Our data involving the role of miR-190 in proper localization of the Scrib/Lgl/Dlg complex could explain the occurrence of miR-190 in several types of cancers. While there are many studies linking miR-190 to a variety of targets in neurogenesis and cancerous outcomes, our understanding of miR-190 as a fundamental principle of cell polarity regulation can explain the many roles of miR-190 in epithelial-mesenchymal transition (EMT), neuronal differentiation and survival, as well as cancer related biological processes

## Methods and Materials

### Fly Strains

The following Drosophila melanogaster stocks were used in this study: w[*]; TI{TI}mir-190[KO]/TM3, P{w[+mC]=GAL4-twi.G}2.3, P{UAS-2xEGFP}AH2.3, Sb[1] Ser[1] (RRID:BDSC_58897) was obtained from BDSC. Oregon-R was used as the control. Flies were maintained on standard yeast-cornmeal-agar medium at 25°C unless otherwise stated. All flies were kept at room temperature in vials containing conventional cornmeal agar medium.

### Embryo collections

Embryos were collected for 0-2 h or 0-4 h after egg laying and aged for 4-5 h at 25°C to get stage 12 embryos. For experiments using mir-190^KO^ (GFP detection), embryos were aged for 4:30 h at 25°C to get stage 12 embryos, and an additional step for miR-190^KO^ strains were to be selected against GFP for homozygosity.

### RNA isolation, reverse transcription, and quantitative reverse transcriptase PCR

RNA was isolated from mir-190^KO^/TM3, GAL4-twi-UAS-2xEGFP, Sb, Ser, and Oregon-R embryos stages 9-12 using the PureLinkTM RNA Mini Kit (InvitrogenTM, 12183025). 75-100 fly embryos were lysed using 1 ml of TRIZOL Reagent, and RNA was extracted according to the manufacturer’s protocol. RNA was eluted in RNAse-free water, and the concentration of each sample was determined. RT-PCR was performed using a T100 Thermal Cycler. The extracted RNA was used for the RT-PCR with amplification by One-Step RT-PCR Kit (LunaScript® Multiplex One-Step RT-PCR Kit, E1555). Agarose gel was made, and PCR samples were run with 100 bp DNA Ladder and TriTrack DNA Loading Dye. (GeneRuler 100 bp DNA Ladder, Thermo Scientific™ SM0241) qRT-PCR was performed using a CFX96 Touch Real-Time PCR Detection System. The extracted RNA was used for the qRT-PCR with an application by SYBR Green PCR master mix (Luna® Universal One-Step RT-qPCR Kit, E3005). Transcript levels were normalized to Actin. The primer pairs for Actin42A are described in Ponton et al., 2011. Primers used in RT-PCR and RT-qPCR are listed as follows:

Pros F-primer 5′-TCACCATCGCCCCTAAAACC-3’ and R-primer 5’-GCTGGAGCAGTGGAGGAAAT-3’; Mira F-primer 5’-CAGCACGACGAGCATGAAAG-3’ and R-primer 5’-GCTCCTCCACCTTTTGACGA-3’; Brat F-primer 5’-CTACTTCTCGGACGTGTGGG-3’ and R-primer 5’-ATCCGTCCCTTGTTGTCCAC-3’; Stau F-primer 5’-AAACGGCAATGAAACCGCTC-3’ and R-primer 5’-TCGCTTTGCCAACTTCTTGC-3’; Numb F-primer 5’-TTTCTCCCATTGCCGAGGTC-3’ and R-primer 5’-GCTGAGCTCCTGACACAGTT-3’; Baz F-primer 5’-GGGCTTTTCGGTCACAACAC-3’ and R-primer 5’-CATTGGAGTGCCATCCACCT-3’; aPKC F-primer 5’-AAGAATCCCGCTGACCGTTT-3’ and R-primer 5’-CGCGATCTGAGTCTAAGCGT-3’; Deadpan 5’-TCAGTTCCGGCCAATCTTCC-3’ and R-primer 5’-TGGGCTTGTGCTGTTCTTCA-3’; Insc F-primer 5’-ACTACCCACAAAGACAGCCAG-3’ and R-primer 5’-GCCAGGATAGCTTGGAGCTG-3’; Lgl F-primer 5’-GCACACTAGACCCACCGAT-3’ and R-primer 5’-CGGTCAGAGCTTAACCGTGT-3’; Dlg F-primer 5’-CTACAGGGTGGAGTTACCCG-3’ and R-primer 5’-CCCAGAATAATCACCGGGCG-3’; Scrib F-primer 5’-CATTGACAAGGATGCCGCAC-3’ and R-primer 5’-TGTCGACGCCAGACTTTGTG-3’; Msi F-primer 5’-GACGTCGTCTGACAAGCTCA-3’ and R-primer 5’-GAGTGTGTGAATGGGCACCT-3’; RhoGAP100F F-primer 5’-GGACGTGACACACATGTCCT-3’and R-primer 5’-CGTCTCGCAAATCTCGCTTG-3’; RhoGAP71E F-primer 5’-AGCGACTGCACAGACTCTTC-3’ and R-primer 5’-CGCTGACGCAGGATTGAAAG-3’; Cdc42 5’-CCCACGGTGTTCGACAACTA-3’ and R-primer 5’-CGAAGGAACTGGGACTGACC-3’.

### Larval Collection

Flies were placed in cages 24 hours prior before first instar larvae collection. Cages were kept at room temperature in dark conditions. For miR-190, homozygous miR-190^KO^ larvae were selected. First instar larvae were selected because homozygous miR-190^KO^ larvae did not survive past 1st instar.

### Brain dissection and dissociation preparation for scRNAseq

After first instar larvae were collected and sorted, larvae were placed in cold PBS on a frozen dissecting glass tray. Fine dissection with forceps was performed on as many larvae as dissected in 45 minutes, any longer time was a concern for cell death and transcriptional changes. 25 control brains were obtained and 35 miR-190^KO^ brains during the 45-minute time period. Brains were collected in tubes with cold Phosphate Buffered Saline (PBS) (Thermo Fisher Scientific AM9624). After collection, tubes were spun down and PBS was removed. 1mL of digestion buffer was added. Digestion buffer included elastase (4mg/ml Worthington Biochemical LS002292), collagenase (2.5 mg/ml Sigma-Aldrich C9722), and cell dissociation buffer (1mL Thermo Fisher Scientific 13-151-014). Tubes were pipetted up and down with a 200ul pipet for 20 minutes. Cells were then filtered through a 50uM filter and then pipetted up and down again for another 10 minutes to reduce clumping. Cells were filtered with a 30uM filter and quenched by adding 500ul of S2 buffer. S2 buffer contains 5mL of Schnider’s insect media (Sigma-Aldrich S9895), 1mL of sterile Fetal Bovine Serum (FBS) (Sigma-Aldrich F0926), and 1.2mL insulin (Sigma-Aldrich I6634). Tubes were then centrifuged for 5 minutes at 4C at 3500 rfc. Supernatant was removed and 1mL of S2 buffer was added. Tubes were again centrifuged for 5 minutes at 4C at 3500 rfc. Supernatant was removed and cells were resuspended with 100ml of resuspension buffer. Resuspension buffer contains 1mL of PBS 0.04% UltraPure BSA (Thermo Fisher Scientific AM2616). After cell concentration was determined using a hemocytometer, cells were diluted to 1000 cells/ml.

### 10x Genomic Sequencing and Seurat data processing

A single-cell suspension concentration of 1000 cells/ml was submitted through UCSF Core Immunology Lab. scRNA-seq libraries were prepared using the Chromium Single Cell 3’ Library and Gel Bead Kit v3 (10X Genomics). The pipeline consists of the following: sample preparation, single cell GEM generation using the 10x Single Cell Controller, sample cleanup, reverse transcription, cDNA preparation, sequencing library preparation, and sample sequencing. Sample preparation involved removal of the dead cells and viability assessment using a flow cytometer. Sequencing was accomplished with HiSeq 4000 or Novaseq.

Seurat version 4.1.0 (Butler et al., 2018; Satija et al., 2015; Stuart et al., 2018) pipeline was adapted and executed on the normal and miR-190 mutant dataset. Quality control (QC) analysis on the control and miR-190^KO^ datasets determined thresholds for filtered mitochondrial counts and unique features. The data was filtered for cells > 10 percent mitochondrial counts and cells that have unique feature counts between 1000 and 25000. A total of 11,580 cells were obtained; 3,951 control cells with a median of 600 genes per cell and 7,629 miR-190^KO^ cells with a median of 1200 genes per cell (Fig. S1). Before integrating, the datasets were preprocessed selecting 2000 features. Next, the datasets were merged through the integration method of SCTransform (SCT). Integration among the datasets was performed by identifying common anchors between the two datasets and combining them into a single Seurat object. To define the dimensionality of the dataset, an ElbowPlot was used to determine [1:20] dimensions. This dimensionality was used to find integration anchors. Using 20 dimensions was also used when applying the “FindNeighbors” function, necessary for clustering. Resolution 0.08 was based on the number of identifiable clusters. UMAP was visualized using the reduction “pca”. Cluster identities were determined based on previously identified gene markers for each cell type.

To determine if there is a difference in the number of cells in each cell type between controls and miR-190^KO^, the number of cells per cluster were counted. Since the number of cells in control and miR-190 mutant is different (3,951 wildtype cells and 7,629 miR-190 mutant cells), the data was normalized by dividing control 3951/ miR-190^KO^ 7629 = 0.518. Each final total cell count of miR-190^KO^ clusters were multiplied by the difference (0.518) before graphing results. Neuronal clusters were sub-clustered forming 8 new clusters. Clusters were visualized in a UMAP plot, with a resolution 0.3. Clusters were identified based on known marker genes.

### Immunofluorescence and confocal imaging

Drosophila embryos were fixed with Heptane saturated with 37% Formaldehyde, as described previously (Rothwell and Sullivan, 2007). The antibodies used were: mouse anti-Prospero MR1A mouse, anti-discs large 4F3 (1:100, Developmental Studies Hybridoma Bank), rabbit anti-Bazooka (1:500 donated by Andreas Wodarz, University of Cologne, Germany), rat anti-Par-6 (1:500), rabbit anti-Scribble (1:500) and rat anti-Lgl (1:100 were donated by Ken Prehoda, University of Oregon), guinea-pig anti-Numb (1:500 donated by Yuh Nung Jan, University of California, San Francisco), rabbit anti-Staufen (1:1000, donated by Daniel St Johnston, Gurdon Institute, University of Cambridge) rat anti-Miranda (1:500, Abcam Cat# ab197788), rabbit Anti-phospho-Histone H3 (1:500 Millipore Cat# 06570) rabbit anti-aPKC (1:500, PKCz (C-20) Santa Cruz, discontinued). To detect primary antibodies, the following Alexa-Fluor conjugated secondary antibodies were used: goat anti-rat Alexa 647 (1:500; Cat# A21247), goat anti-rabbit Alexa 647 (1:500; Cat# A21244), goat anti-rabbit Alexa 555 (1:500; Cat# A21429), goat anti-guinea-pig Alexa 555 (1:500; Cat# A21435), goat anti-mouse Alexa 488 (1:500; Cat# A11001), DAPI for nucleic acid staining (1:500; Millipore Cat# D9542).

First instar larvae were collected, sorted and placed in PBS on a dissecting glass tray. Dissection of larval brains was performed, and brains were placed into a 1.5mL tube with 1x PBS 0.01% Triton X-100. Once the brains were collected, the supernatant was removed and 1mL of 4% paraformaldehyde (PFA) was added. Tubes were incubated on a rotator at room temperature for 30 minutes. After 30 minutes, PFA was removed, and brains were washed with 1mL of PBST three times for 10 minutes. Brains were blocked in 1mL goat buffer for 1 hour. After the hour, the buffer was removed and 1mL PBST was added. The following primary antibodies and their concentrations were used: anti-Prospero [1:1000], repo [1:1000]. Brains were incubated with primary antibodies overnight on a rotator at 4C. After incubation, brains were washed three times with PBST for 10 minutes. Secondary antibody was added in 1mL PBST and incubated at room temperature for 1 hour. Brains were washed three times with PBST before being fine dissected and mounted on a slide using glycerol mounting media.

Microscopy was performed using a Zeiss LSM 710 and LSM 980; Leica Stellaris image processing and quantification were performed with the open-source software FIJI. To calculate the average number of cells per lobe, each larval lobe had five z-stacks images performed. All cells of interest in each z-stack image were counted and added together. The sum of cells in each lobe was averaged per sample and compared.

### Quantification and Statistical Analyses

All microscopy data represent images from at least three biological replicates (n), as specified in the figure legends. For each replicate, z-stacks were acquired, processed, and analyzed using ImageJ (Schindelin et al., 2012). Quantification graphs display mean ± standard deviation as indicated in the respective figure legends, with statistical significance determined by an unpaired t-test. All statistical analyses were conducted using Prism (GraphPad), and p-values are provided in charts or figure legends as indicated.

## Data availability statement

All data supporting the findings of this study are available within the article and its supplementary materials. Raw and processed datasets have been deposited in [Name of repository] under accession number [Accession number]. Additional data that support the conclusions of this study are available from the corresponding author upon reasonable request.

All custom code used for data analysis is available at [GitHub/Zenodo link, if applicable].

## Acknowledgments

We would like to thank Todd Nystul, Scott Roy, and Matt de Cruz for technical support and advice on this project. We would also like to thank Annette Chan and the Cell and Molecular Imaging Center (CMIC) at San Francisco State University (SFSU) for technical expertise and use of imaging equipment. L.G., A.N., C.T., and B.R. was funded by a National Science Foundation (NSF) Facilitating Research at Predominantly Undergraduate Institutions award 2127729, NSF Science and Technology Center – Center for Cellular Construction (CCC) DBI-1548297, National Institute of Health (NIH) Bridge to Doctorate T32-GM142515, NIH U-RISE T34-GM145400. L.M.G. was supported by NIDDK K01DK132488, a Koret Foundation Catalyst Award, and a Baxter Foundation Faculty Scholar Award. G.A. was supported by an NSF Graduate Research Fellowship DGE-2146755. This research was supported in part by the PAIR-UP Imaging Science Program which is funded by grants from the Gordon and Betty Moore Foundation, Burroughs Wellcome Fund and CZI.

## Author contributions

G. Ascencio performed experiments, data analysis, figure preparation, writing of the manuscript, review and editing. L. Galvan performed experiments, data analysis, figure preparation, writing and editing. J. Sanchez performed data analysis and figure preparation, writing, and editing. A. Nagainis performed experiments and data analysis. C. Tam performed experiments and data analysis. L. Goins performed data analysis, review and editing. B. Riggs performed planning and methodology, data analysis, writing of the manuscript, review and editing.

## Abbreviations

CNS: Central Nervous System
NB: Neuroblast
scRNAseq: single cell RNA sequencing
ACD: Asymmetric cell division
miRNAs: microRNAs
DEG: differential gene expression

## Supplemental material

### Supplemental Figures

**Fig. S1. QC analysis.** (A-B) Quality control (QC) analysis using Seurat R package of the combined Seurat object containing both the control and miR-190^KO^ conditions that determined thresholds for filtered mitochondrial counts and unique features. (B) ElbowPlot determining dimensionality for UMAP. ElbowPlot shows justification for chosen [1:29] dimensionality.

**Fig. S2. Uniform Manifold Approximation and Projection (UMAP) of Control and miR-190^KO^ First Instar Larvae**. (A-B) Uniform Manifold Approximation and Projection (UMAP) of the integrated dataset separated by experimental condition. Each dot represents an individual cell, with cell clusters annotated based on the expression of known marker genes.

**Fig S3. Quantification of Cell Proportions and Counts of Annotated Cell Types Between Experimental Conditions**. (A) Bar chart showcasing the proportion of the annotated cell types in control and miR-190^KO^ conditions. Bar chart showing the (B) normalized cell count and (C) proportional ratio of the annotated cell types across both experimental conditions.

**Fig. S4. Quantification of GMC and NB population in miR-190 mutants.** To further validate the changes occurring in miR-190^KO^ mutants, we quantified both variables. A) Side by side comparison of Drosophila first instar larval brain staining with Prospero (green), Deadpan (red), and DAPI (blue) between wildtype and miR-190KO mutant. The mean difference between control and miR-190 mutant was 21.79 within the 95% confidence interval of 16.77 to 26.80 p-value being less than 0.0001. B) In miR-190 mutants, there was a significant elevation of GMC population by performing a cell count of Prospero staining; control averaged at 15 cells and mutant averaged at approx. 36 cells. The mean difference between wildtype and miR-190 mutant was-19.00 within the 95% confidence interval of-26.45 to-11.55 making the p-value being 0.0010. C) In contrast, we performed a similar cell count method and observed that neuroblast conversely decreased in mutants compared to wildtype via counting deadpan staining; controls averaged at 40 cells and mutant averaged 20 cells. The mean difference between wildtype and miR-190 mutant was 38.73 within the 95% confidence interval of 14.04 to 63.43 p-value being 0.0071.

**Fig. S5. Spindle rotation defect in *miR-190* knockout (KO) embryos** miR-190KO embryos were fixed and immunostained for Numb (magenta) and DAPI (DNA, cyan). In mitotic neuroblasts (NBs) lacking miR-190, Numb is mislocalized along the cortex, while DNA exhibits a spindle rotation defect. Scale bar: ∼5 μm.

**Supplemental Table 1. List of Marker Genes per Cluster.** Marker genes for each cluster in the integrated dataset were identified using default parameters for the FindAllMarkers function (min.pct=0.25, only. pos=TRUE).

**Supplemental Table 2. List of Differential Expressed Genes (DEGs) in miR-190^KO^ larval brains.** A list of 633 total genes were identified as DEGs, with an adjusted p-value < 0.05 and absolute log2 fold change (log2FC) > 0.58.

**Supplemental Table 3. List of Gene Ontology Terms.** Gene Ontology (GO) terms, showing the number of genes enriched in each term. DEGs were defined by an adjusted p-value < 0.05 and an average log2FC > 0.58. GeneRatio represents the proportion of genes in a term relative to the total number of genes in the analysis.

**Supplemental Table 4. List of Notable miR-190 mRNA targets.** Using TargetscanFly prediction software, 331 potential targets were identified for miR-190-5p and 498 targets for miR-190-3p. Targets included on this list were selected based on an aggregate PCT score of 0.90 or above indicating a less than 10% probability of being conserved by chance. Additionally, genes that displayed a role in neural development and function, cell polarity, and cell fate were included. Conserved miRNA sites include 8mer, 7mer-m8, and 7mer-A1 identifying the number and location of the binding. Key words and gene descriptions were provided by Flybase.

