## Supplementary figures and images for "miR-190 is a Key Regulator in Establishing Cell Polarity and Specification in the Drosophila Nervous System"

### Fig S1

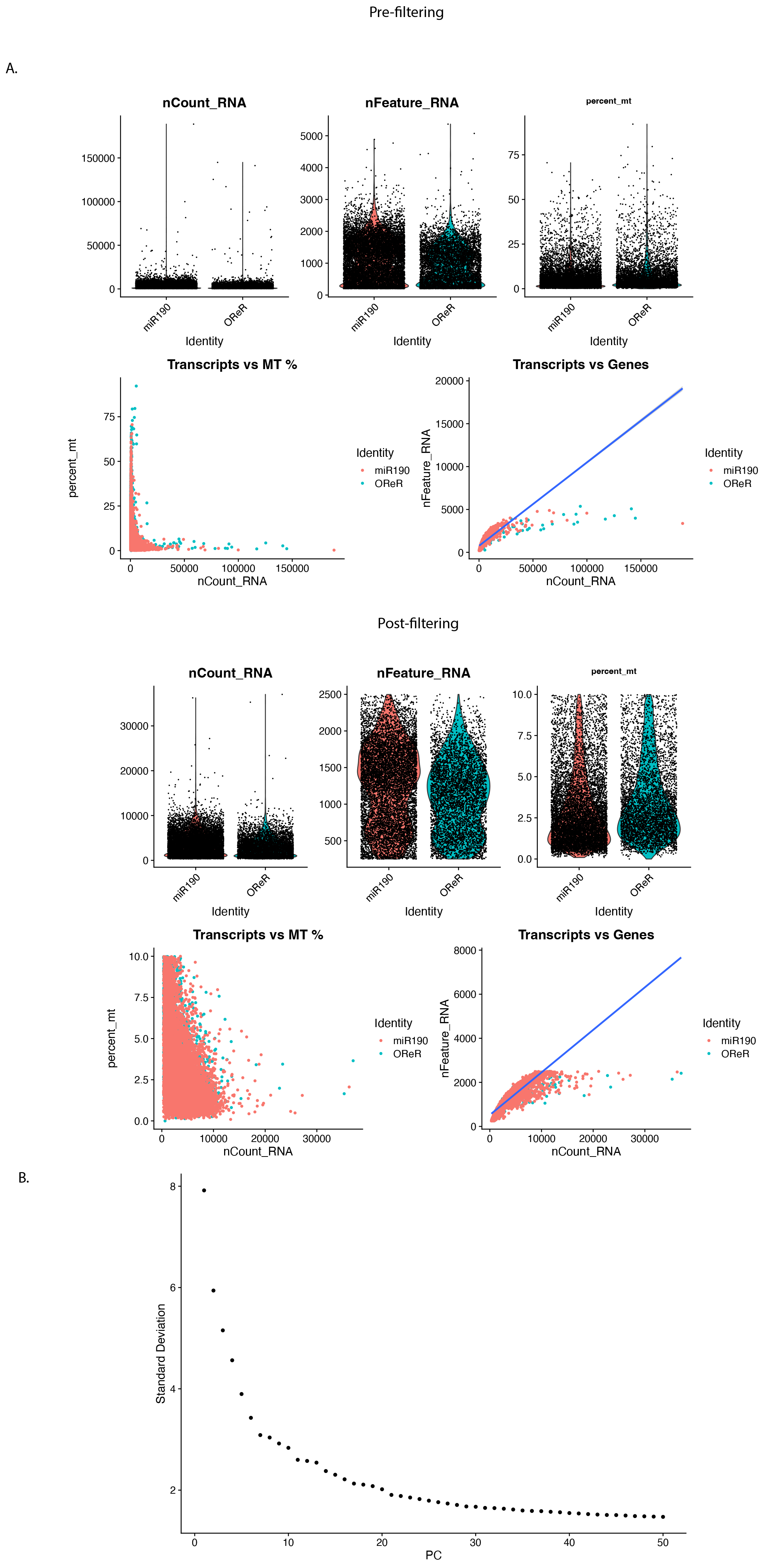

### Fig S2

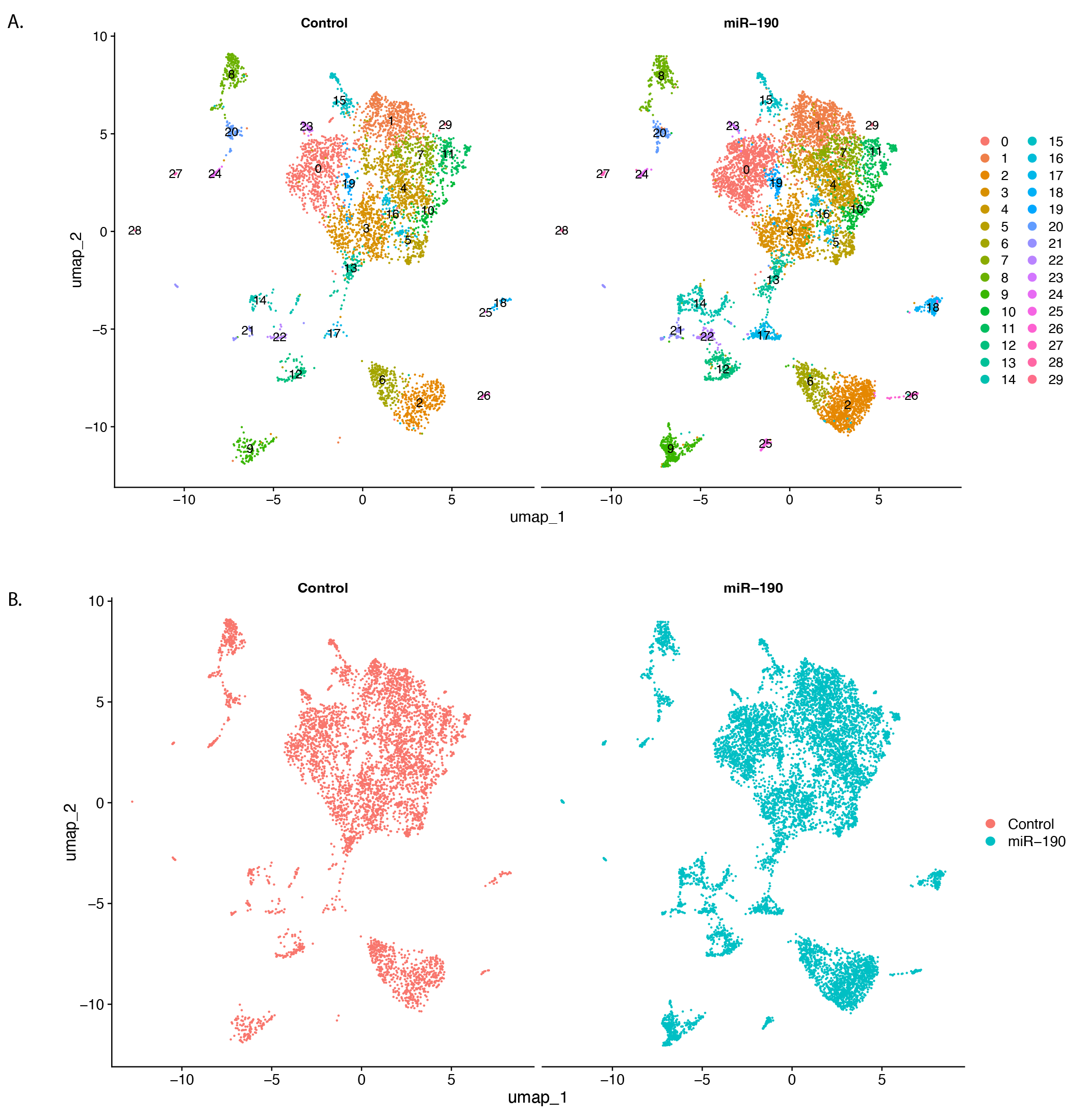

### Fig S3

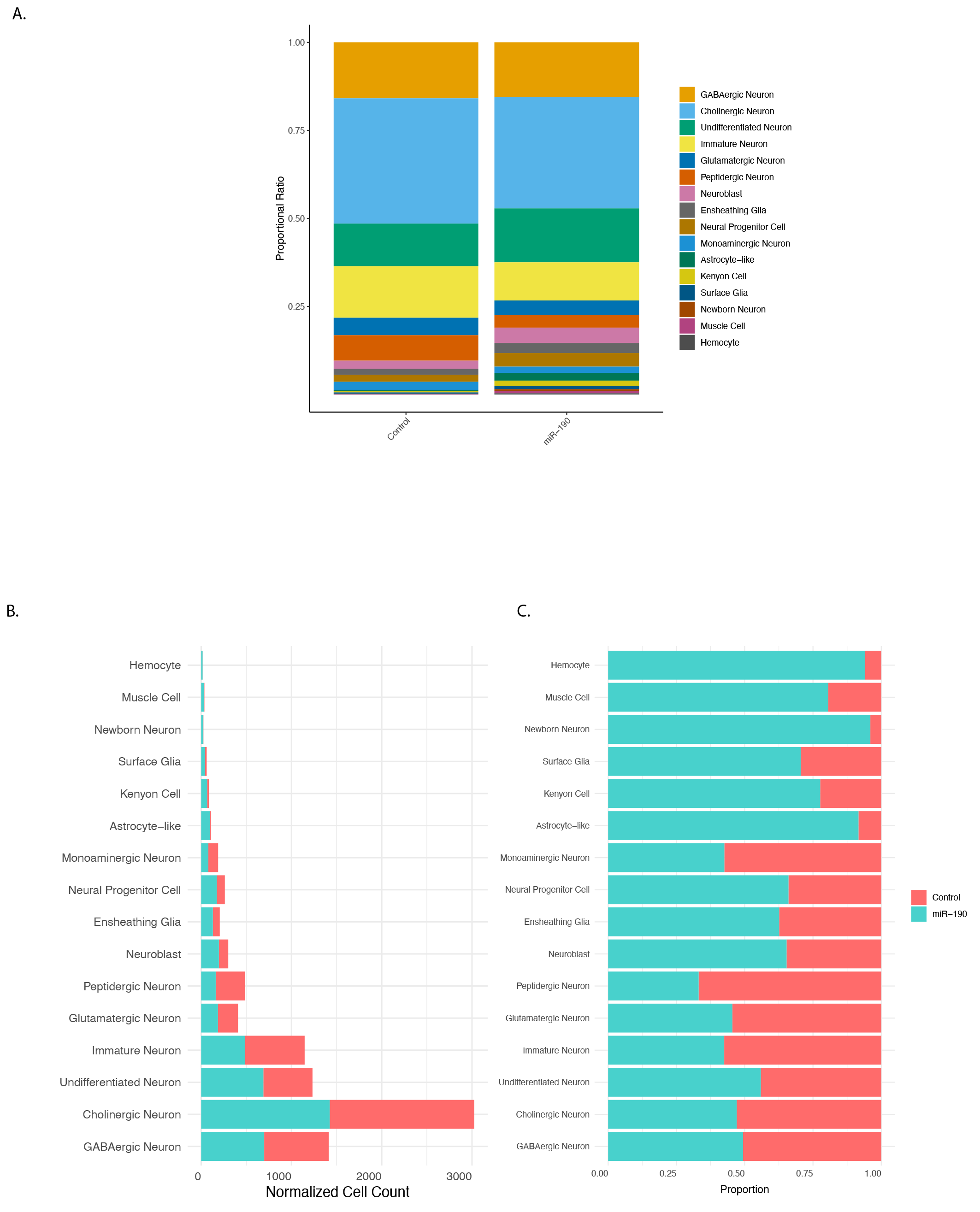

### Fig S4

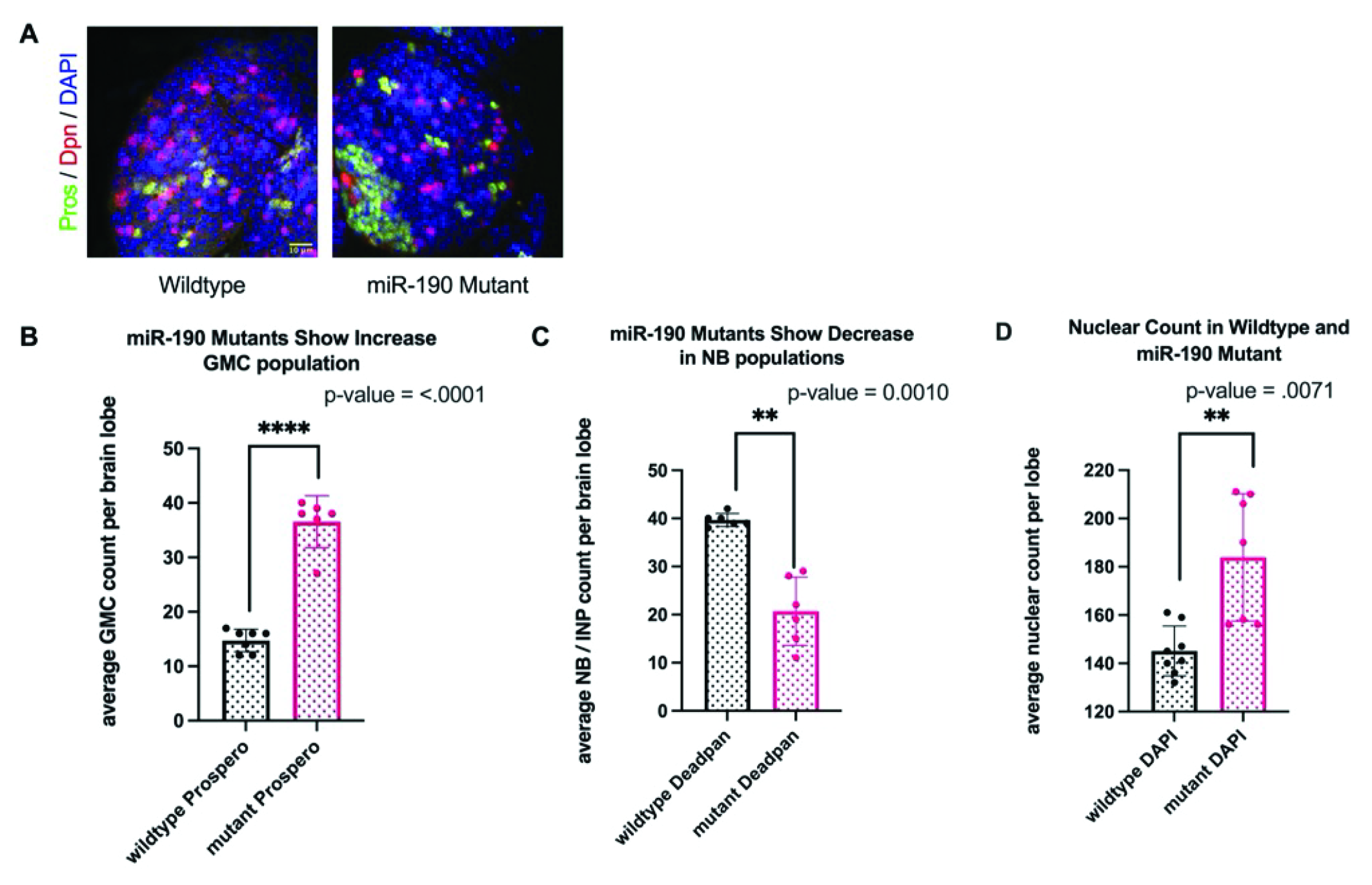

### Fig S5

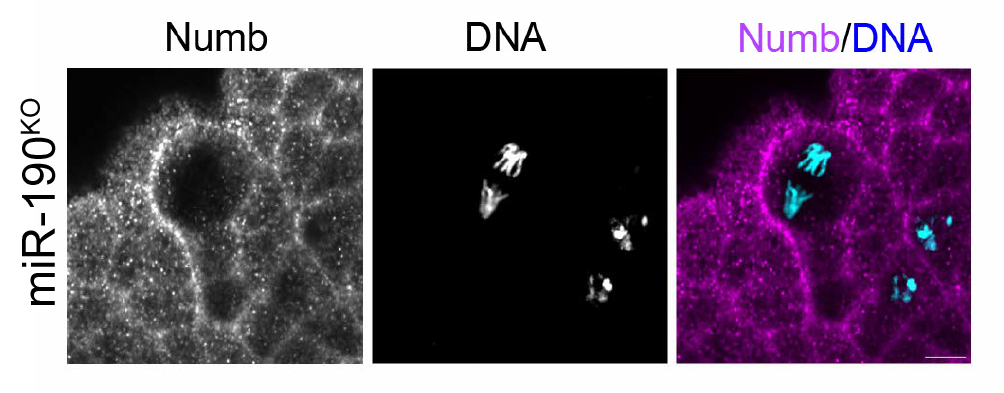
